# Rapid inactivation of vaccinia, a surrogate virus for monkeypox and smallpox, using ultraviolet-C disinfection

**DOI:** 10.1101/2022.10.04.510918

**Authors:** Carolina Koutras, Richard L. Wade

## Abstract

Human monkeypox is an emerging health threat that has the potential to cause serious sequelae. Ultraviolet-C (UV-C) disinfection is a physical process that triggers microbial inactivation through irreversible DNA damage. A high-output mobile UV-C unit was evaluated against vaccinia, a monkeypox surrogate, for antimicrobial efficacy. In under 7 minutes, a single UV-C cycle had a virucidal efficacy of ≥ 99.996 % in a 200 sq feet area. UV-C technology is a promising strategy for infection prevention and control in the post-COVID era.

## Background

Human monkeypox is a rare viral zoonosis caused by the monkeypox virus belonging to the genus *Orthopoxvirus* of the *Poxviridae* family. Endemic to central and western Africa, human monkeypox has recently emerged in the USA and was declared a public health emergency of international concern. *Poxviridae* are a diverse group of large double-stranded DNA viruses that replicate exclusively in the cytoplasm of infected cells. The most well-known member of the *Orthopoxvirus* genus is the now extinct variola virus, the causative agent of smallpox. Other notable members of the genus include the vaccinia virus, which is used in the current smallpox vaccine, and cowpox virus. These viruses are similar in terms of size, shape, replication, and structure. With few exceptions, they fail to trigger a chain of transmission in humans and remain zoonotic. However, outbreaks of human monkeypox have emerged in the last 50 years and are of concern for several reasons. There is currently no proven treatment for human monkeypox; clinical efficacy data on the use of smallpox vaccines against monkeypox is lacking; the virulence and transmissibility of monkeypox in humans is not fully understood; the disease has the potential to result in major disease sequelae, including disfiguring scars and permanent corneal lesions; and finally, monkeypox virions can persist in the environment for long periods of time. ^1-4^ The goal of this study was to evaluate the efficacy of an ultraviolet-C (UV-C) whole-room disinfection tower against monkeypox on contact surfaces, using the highly similar vaccinia virus as a surrogate.

## Methods

Three replicates of glass test and control carriers were inoculated with a 0.2 ml volume of viral suspension (modified vaccinia virus Ankara strain, ATCC VR-1508) and dried at ambient room temperature (19.2-19.4°C) and 48% relative humidity (RH). A 78” tall UV-C device equipped with 8 high-output lamps and reflectors, and 4 long-range passive infrared safety sensors (Arc, R-Zero Systems) was placed at 8 feet (∼200 sq feet radius) from the test carriers. Carriers were exposed to UV-C light for 6 minutes and 52 seconds in total. Following harvest of test and control carriers, the viral suspensions were quantified using the TCID_50_ (Median Tissue Culture Infectivity Dose) technique. The inoculated cell culture plates were incubated for a few days and microscopically scored for the presence/absence of the test virus. The Spearman-Kärber method was used for estimating viral titers. The log10 and percent reductions in viral titer were calculated for UV-C exposed carriers relative to controls. The study diagram is shown in Figure 1.

**Figure 1.**
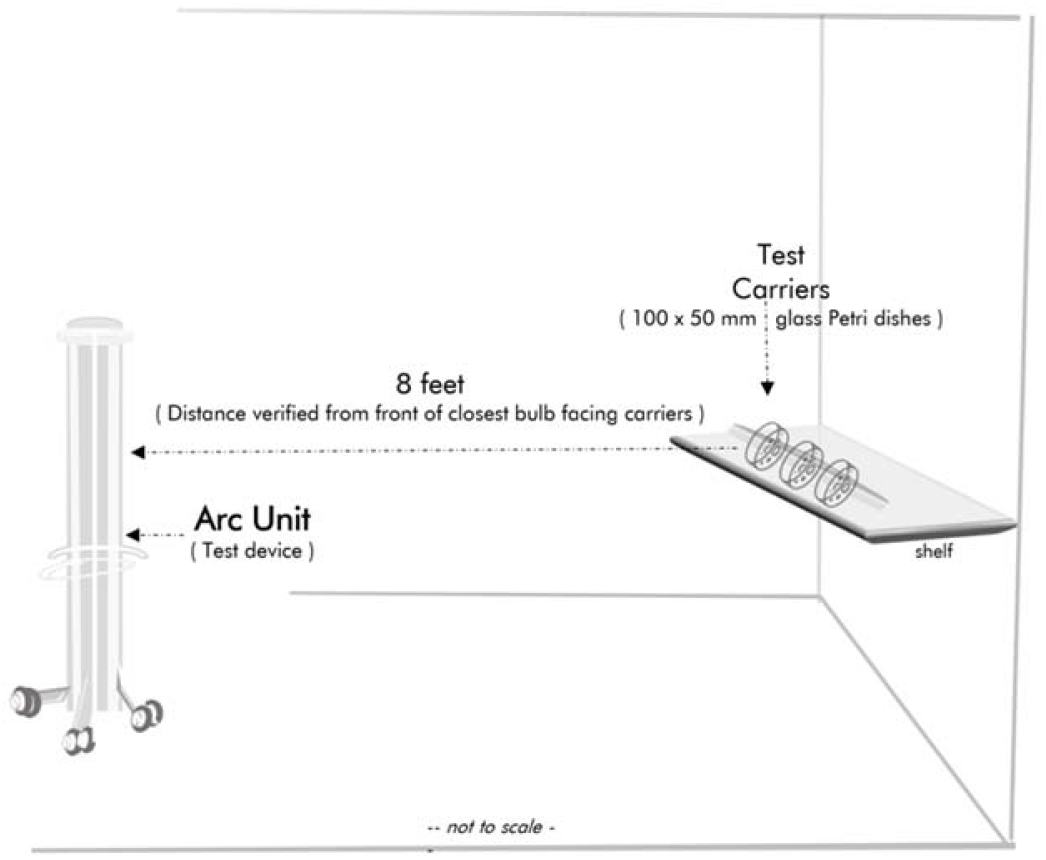
Study Diagram. The test device was placed at an 8-feet distance from the test carriers.

## Results

Table 1 summarizes the TCID_50_ per carrier, and average TCID_50_ for both test and control conditions. The recovery control plate had an average viral titer of 5.30 log10 TCID_50_ per carrier. The test plate corresponding to the UV-C disinfection treatment had a ≥ 4.40 log10 reduction (≥ 99.996%) in viral titer relative to the titer of the corresponding recovery control plate.

**Table 1.**
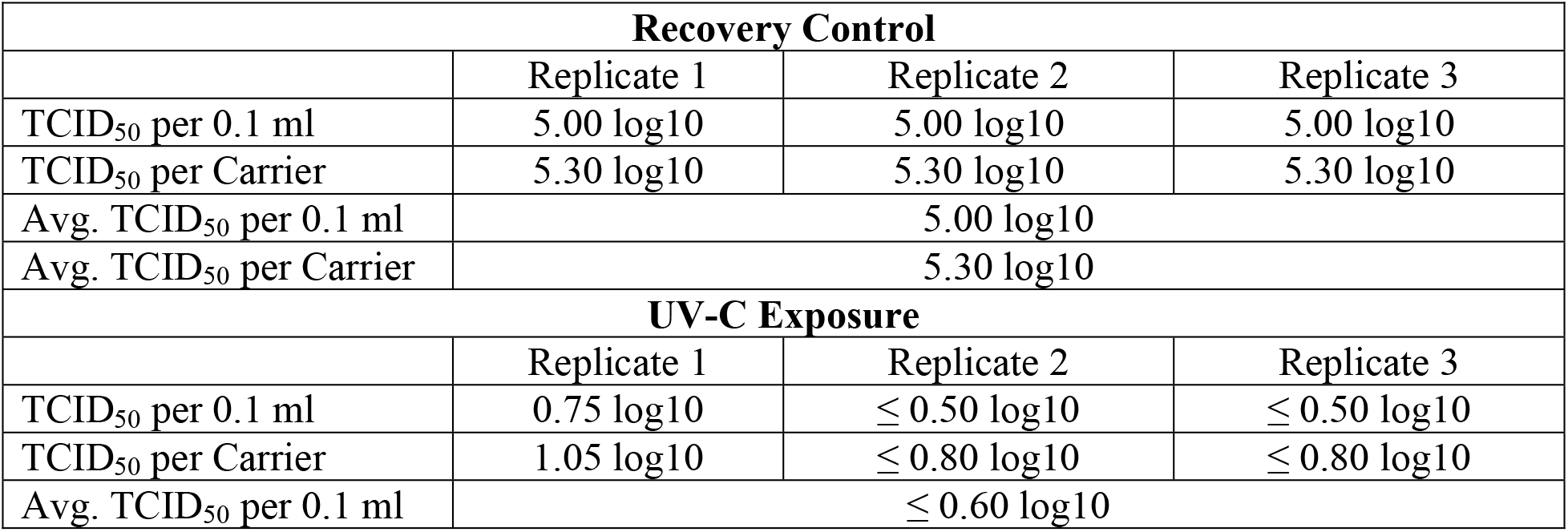
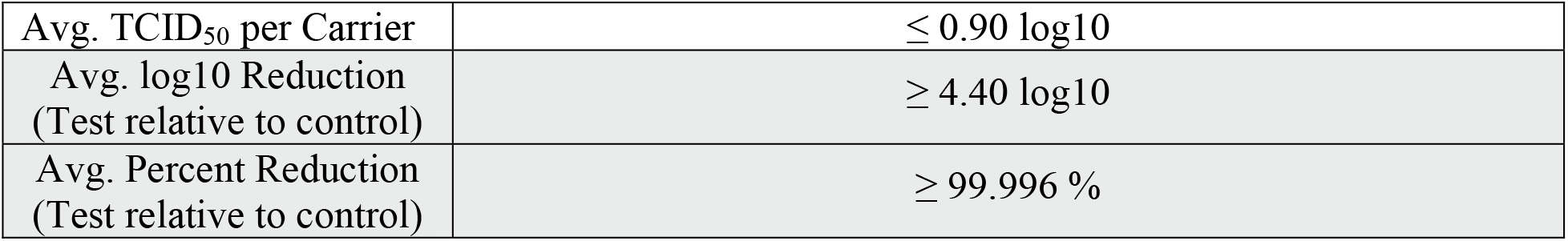
The average TCDI_50_ per carrier is shown for both the recovery and test replicates. The average log10 and percent reduction for the test condition relative to control is also provided.

| Recovery Control |  |  |  |
| --- | --- | --- | --- |
|  | Replicate 1 | Replicate 2 | Replicate 3 |
| TCID<sub>50</sub> per 0.1 ml | 5.00 log10 | 5.00 log10 | 5.00 log10 |
| TCID<sub>50</sub> per Carrier | 5.30 log10 | 5.30 log10 | 5.30 log10 |
| Avg. TCID<sub>50</sub> per 0.1 ml | 5.00 log10 |  |  |
| Avg. TCID<sub>50</sub> per Carrier | 5.30 log10 |  |  |
| UV-C Exposure |  |  |  |
|  | Replicate 1 | Replicate 2 | Replicate 3 |
| TCID<sub>50</sub> per 0.1 ml | 0.75 log10 | ≤ 0.50 log10 | ≤ 0.50 log10 |
| TCID<sub>50</sub> per Carrier | 1.05 log10 | ≤ 0.80 log10 | ≤ 0.80 log10 |
| Avg. TCID<sub>50</sub> per 0.1 ml | ≤ 0.60 log10 |  |  |

Table 1. The average TCID<sub>50</sub> per carrier is shown for both the recovery and test replicates. The average log<sub>10</sub> and percent reduction for the test condition relative to control is also provided.
|  |  |
| --- | --- |
| Avg. TCID<sub>50</sub> per Carrier | $\leq 0.90 \log_{10}$ |
| Avg. log<sub>10</sub> Reduction<br>(Test relative to control) | $\geq 4.40 \log_{10}$ |
| Avg. Percent Reduction<br>(Test relative to control) | $\geq 99.996 \%$ |

## Discussion

Monkeypox is an important emerging pathogen that can lead to deep seeded lesions across the body and may result in more infections than originally believed. In this study, a mobile UV-C tower equipped with high-performance bulbs successfully inactivated ≥ 99.996 % (≥ 4.40 log10) of the vaccinia virus, a monkeypox surrogate, in less than 7 minutes. There are two published reports exploring the susceptibility of vaccinia aerosols to UV-C. ^5,6^ However, prolonged direct contact with a virion source is the emerging transmission route for human monkeypox. Technological innovations in recent years led to the development of faster, accessible, and high-performing UV-C devices ^7^ that also incorporate safety sensors to eliminate accidental UV-C exposures. While this study focused on an emerging pathogen, the germicidal properties of UV-C light are not species-specific. Rather, UV-C inactivates microorganisms by causing damage to their nucleic acids. In conclusion, UV-C disinfection is an effective pathogen suppression technology that can be easily deployed in healthcare and community settings for the prevention and control of pathogens that may persist in the environment.

## Acknowledgements

The authors would like to thank Dr. Benjamin Tanner and Microchem Laboratories (Austin, TX) for their commitment to this project.

